# Suppression of *Plasmodium* MIF-CD74 Signaling Protects Against Severe Malaria

**DOI:** 10.1101/2021.02.14.430970

**Authors:** Alvaro Baeza Garcia, Edwin Siu, Xin Du, Lin Leng, Blandine Franke-Fayard, Chris J Janse, Shanshan W Howland, Laurent Rénia, Elias Lolis, Richard Bucala

**Affiliations:** Department of Internal Medicine, Yale School of Public Health, New Haven, Connecticut 06520, USA; Departments of Pharmacology, Yale School of Medicine and Epidemiology of Microbial Diseases, Yale School of Public Health, New Haven, Connecticut 06520, USA; Department of Parasitology, Leiden University Medical Center, Leiden, Netherlands; Singapore Immunology Network, Agency for Science, Technology and Research (A*STAR), Singapore, Singapore; Department of Pharmacology, School of Medicine, Yale University, New Haven, CT, 06510, USA

## Abstract

Malaria begins when mosquito-borne *Plasmodium* sporozoites invade hepatocytes and usurp host pathways to support the differentiation and multiplication of erythrocyte-infective merozoite progeny. The deadliest complication of infection, cerebral malaria, accounts for the majority of malarial fatalities. Although our understanding of the cellular and molecular mechanisms underlying the pathology remains incomplete, recent studies support the contribution of systemic and neuroinflammation as the cause of cerebral edema and blood-brain barrier (BBB) dysfunction. All *Plasmodium* species encode an orthologue of the innate cytokine, Macrophage Migration Inhibitory Factor (MIF), which functions in mammalian biology to regulate innate responses. *Plasmodium* MIF (PMIF) similarly signals through the host MIF receptor CD74, leading to an enhanced inflammatory response. We investigated the PMIF-CD74 interaction in the onset of experimental cerebral malaria (ECM) using CD74 deficient (*Cd74*−/−) mice, which were found to be protected from ECM. The protection was associated with the inability of brain microvessels from *Cd74*^−/−^ hosts to present parasite antigen to sequestered *Plasmodium*-specific CD8^+^ T cells. Infection of mice with PMIF-deficient sporozoites (*Pb*A*mif*-) also protected mice from ECM, highlighting the pivotal role of PMIF in the pre-erythrocytic stage of the infection. A novel pharmacologic PMIF-selective antagonist reduced PMIF/CD74 signaling and fully protected mice from ECM. These findings reveal a conserved mechanism for *Plasmodium* usurpation of host CD74 signaling and suggest a tractable approach for new pharmacologic intervention.

## Introduction

Malaria caused by parasites of the genus *Plasmodium* is the most deadly parasitic disease, causing approximately half a million deaths annually [1]. *Plasmodium* sporozoites enter the skin through the bite of infected *Anopheles* mosquitoes and transit through the bloodstream to invade the liver. A single infected hepatocyte produces tens of thousands of erythrocyte-infectious merozoites and initiates the erythrocytic cycle of infection. The subsequent erythrocytic stage of infection produces the disease’s clinical manifestations [2], including the most severe complication of *P. falciparum* infection: cerebral malaria leading to impaired consciousness, seizures, coma, and subsequent mortality [3]. The experimental cerebral malaria (ECM) animal model by infection of susceptible C57BL/6J mice with *Plasmodium berghei* ANKA (*Pb*A) reproduces many of the neurological signs and pathologic changes of human cerebral malaria [4]. ECM is triggered by parasitized erythrocytes in the cerebral microvasculature leading to the production of inflammatory molecules such as IFN-γ, granzyme B, and perforin, and is associated with the recruitment and accumulation of effector CD8+ T cells in the CNS [5, 6].

Both host and parasite factors contribute to the pre- and erythrocytic stages of infection and severe malaria development. *Plasmodium* parasites express intricate strategies to evade immune detection and destruction. It is noteworthy that all *Plasmodium* species analyzed genetically encode an orthologue of the mammalian cytokine macrophage migration inhibitory factor (MIF) [7, 8]. MIF sustains activation responses by promoting innate cell survival, which occurs by signaling through its cognate receptor CD74, leading to sustained ERK1/2 activation and reducing cellular p53 activity [9, 10, 11]. *Plasmodium* MIF (PMIF) is highly conserved in all known *Plasmodium* genomes; for instance, only a single amino acid distinguishes murine *Plasmodium berghei* from human *P. falciparum* PMIF [7, 8]. Recent evidence has implicated PMIF in the growth and development of liver-stage parasites [12, 13], and PMIF binds with high affinity to the host receptor CD74 [14, 15], which has been independently identified as a susceptibility factor for murine *Plasmodium* infection [16].

In the present study, we show that *Pb*A infected *Cd74*−/− mice are resistant to ECM. Cerebral malaria onset further relies on the contribution of endothelial cell CD74, which is upregulated in the brains of infected mice in the presence of PMIF to promote parasite antigen presentation to brain-sequestered *Plasmodium*-specific CD8^+^ T cells. Mice infected with *PbAmif*- parasites were only resistant to ECM when infected with sporozoites, reinforcing the idea that liver-stage *Plasmodium* infection is critical for ECM development [17]. In agreement with prior studies [12, 13], our data also support a central role for PMIF in *Plasmodium* liver infection. PMIF activates the hepatocellular host MIF receptor CD74 to inhibit the apoptosis of infected hepatocytes, thus promoting *Plasmodium* development and replication. These findings were recapitulated by pharmacologic inhibition of the PMIF/CD74 interaction with a novel, PMIF-selective small molecule antagonist [18, 19] that reduced the survival of infected cells, decreased liver-stage parasite burden, and fully protected mice from acute cerebral malaria.

## Results

### CD74 is overexpressed in the brain of *Pb*A infected mice and contributes to ECM development

We recently demonstrated that PMIF exerts its proinflammatory effects by signaling through the host receptor CD74 [20]. To examine the potential role of CD74 in the pathogenesis of ECM, we measured the expression of CD74 in *Pb*A-infected mouse brains during ECM and observed an increase in *Cd74* mRNA expression compared with uninfected mice (**Figure 1A**). We next challenged WT and *Cd74*^−/−^ mice with *Pb*AWT iRBCs and assessed ECM development. While 100% of the WT mice exhibited neurological symptoms within 7-8 days after infection, *Pb*A-infected *Cd74*^−/−^ mice were fully protected from ECM and succumbed to hyperparasitemia only 30 days after infection (**Figure 1B, C and S1A**). Moreover, we found no significant differences in parasitemia between *Pb*AWT-infected WT or *Cd74*^−/−^ mice during the asymptomatic blood-stage, suggesting that *Cd74* deficiency does not affect parasite replication in the erythrocyte (**Figure S1B**). The same results were observed in *Cd74*^−/−^ mice infected with *Pb*A*mif*- parasites (**Figures S1C, D, and E**). The protection of *Cd74*^−/−^ mice was associated with the downregulation of IFN-γ, perforin, and granzyme B expression in the brains of *Cd74*^−/−^ versus WT mice but without an appreciable difference in the quantity of brain sequestered parasites (**Figure 1D and S1F)**.

**Figure 1.**
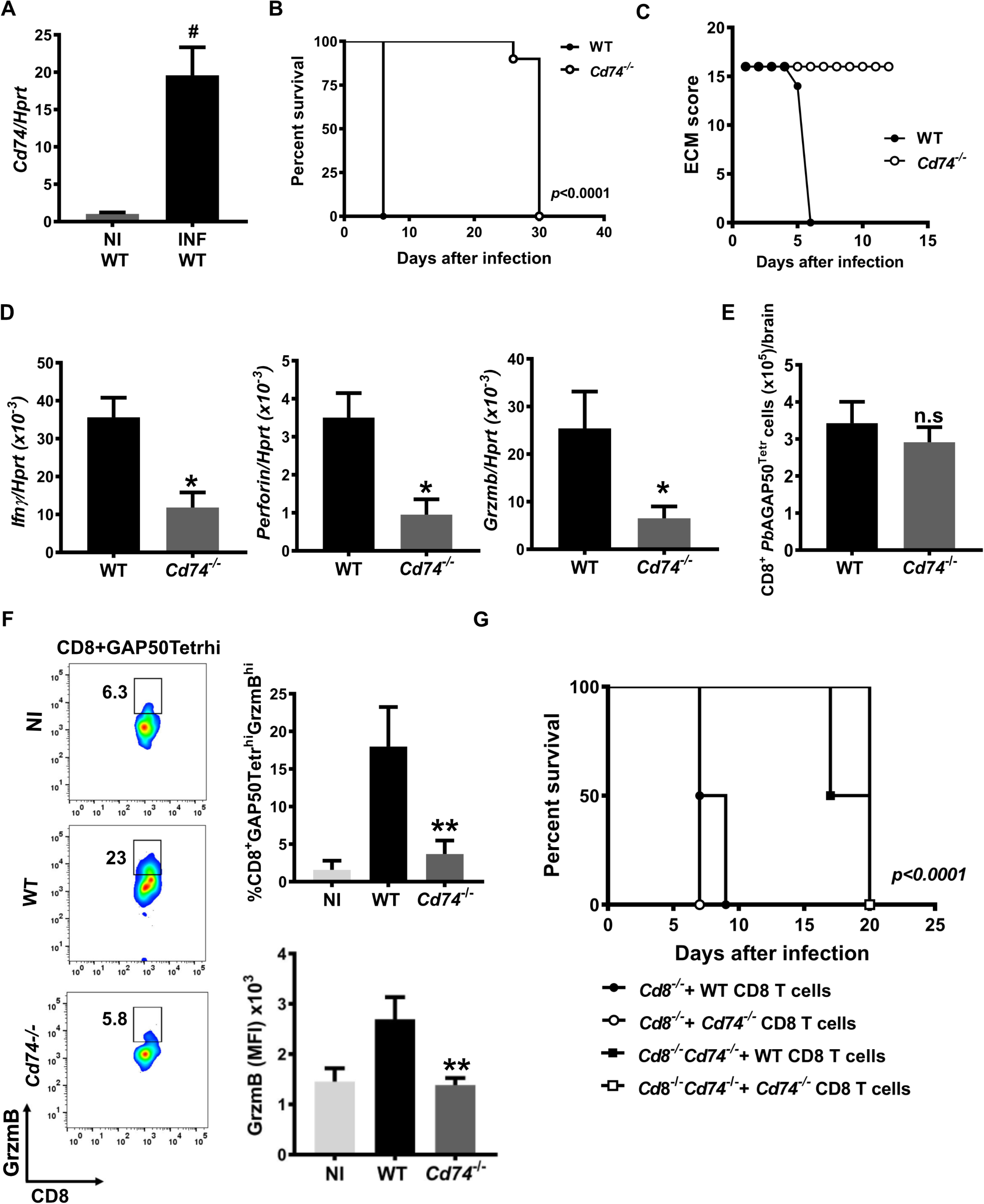
CD74 is overexpressed in the brain of *Pb*A infected mice and contributes to ECM development. **A,***Cd7*4 mRNA expression measured by qPCR in the brains of wild type (WT) C57BL6/J mice that were either non-infected (NI WT) or infected with 10^6^ *Pb*A iRBC (INF WT). Results are shown as mean ± SD from two independent experiments with 6 animals per group and experiment: #p<0.0001 by Mann-Whitney test. CD74 deficient (*Cd74^−/−^*) and WT C57BL6/J mice were infected i.p with 10^6^ *Pb*A iRBC and the **B**, Kaplan–Meier survival plots for WT and *Cd74*^−/−^ mice following infection with *Pb*A and **C,** ECM malaria score were assessed. Data are from two pooled independent experiments with 10 animals per group; p < 0.0001 by log-rank (Mantel Cox) test. Bars represent the ± SD of n=10 WT and n=10 *Cd74*^−/−^ mice and pooled from three independent experiments. **D**, Transcriptional expression of IFNγ, perforin and granzyme B was measured in brain tissue of PbA infected WT and Cd74−/− mice on day 7 after infection by quantitative real-time PCR. Results are expressed as mean ± SD of n=6 mice per group pooled from two experiments: *p=0.0159 and *p=0.0317 by two-tailed Mann-Whitney test. Brain infiltrating lymphocytes from WT and *Cd74^−/−^ Pb*A infected mice were isolated 7 days after infection, and the number of **E**, pathogenic tetramer-labeled brain CD8+ T cells (CD8^+^GAP50Tetra^hi^), expressing **F**, the proinflammatory marker GrzmB (CD8^+^GAP50Tetra^hi^GrzmB^hi^) measured by flow cytometry. Data are shown as mean ± SD of n=6 WT and n=6 *Cd74*^−/−^ mice and pooled from two independent experiments; n.s.=non-significant; *p=0.0022 by two-tailed Mann-Whitney test. **G,** Kaplan-Meier survival plots for C57BL6/J mice *Cd8*^−/−^ and *Cd8*^−/−^*Cd74*^−/−^ receiving CD8^+^ T cells isolated from *Pb*A infected WT or *Cd74*^−/−^ and infected with i.p with 10^6^ *Pb*AWT iRBC. Data are form three pooled independent experiments with 6 animals per group; p<0.0001 by log-rank (Mantel Cox) test.

CD8^+^ T cells are essential for ECM development and contribute directly to human cerebral malaria [4] and ECM pathology [21]. Thus, we investigated if CD8^+^ T cells from *Cd74*^−/−^ mice have an impaired response to *Pb*A infection. We measured the amount of brain sequestered CD8^+^ T cells responding to *Pb*A by using a T cell receptor tetramer specific to the *Pb*AGAP50 antigen [22]. Notably, the amount of brain-sequestered *Pb*AGAP50-specific CD8^+^ T cells was not significantly different between WT and *Cd74*^−/−^ mice (**Figure 1E**), indicating that *Cd74*^−/−^ mice can mount a *Pb*A responsive CD8^+^ T cells response in the brain. Nevertheless, in *Cd74*^−/−^ mice, CD8 T cell effector functions were strongly suppressed, as indicated by the reduced frequency of *Pb*GAP50-specific CD8^+^ T cells expressing the ECM-associated inflammatory molecule Granzyme B (**Figure 1F**). Our findings suggest that *Pb*A responsive CD8^+^ T cells from *Cd74^−/−^ mice* undergo priming and trafficking to the brain during *Pb*A blood-stage infection but do not express the inflammatory effector response associated with the development of ECM. We therefore examined if a dysfunctional cytotoxic response in *Cd74* deficient CD8^+^ T cells reduced ECM symptoms. For this, we adoptively transferred CD8^+^ T cells from WT or *Cd74^−/−^ Pb*A-infected mice into naïve *Cd8*^−/−^ or *Cd8*^−/−^*Cd74*^−/−^ recipient mice and infected them with *Pb*A-infected red blood cells (iRBCs) three days later. Recipient *Cd8*^−/−^ mice that received WT or *Cd74*^−/−^ CD8^+^ T cells from *Pb*A-infected mice showed signs of ECM and succumbed by day 10, whereas recipient *Cd8*^−/−^ *Cd74*^−/−^ mice receiving WT or *Cd74*^−/−^ CD8^+^ T cells from *Pb*AWT infected mice did not show ECM symptoms and succumbed by 20 days after infection (**Figure 1G**). Together, these data support the conclusion that CD8^+^ T cells from C*d74*^−/−^ mice are primed by *Pb*A antigens but are unable to induce ECM, supporting the role of brain expressed CD74 in the development of ECM pathology.

### Cross-presentation of *Plasmodium* antigen by brain endothelium is CD74 dependent

Brain vascular endothelium becomes activated during malaria infection with the ability to process and cross-present *Plasmodium* antigens [22], thereby contributing to the T cell effector response and inflammation that underlies ECM [21]. In addition to the role of CD74 as the cognate MIF receptor [10], it functions intracellularly as the MHC class II invariant chain [23] and has been implicated in an MHC class I cross-presentation pathway for cytolytic T lymphocytes (CTL) [24]. We hypothesized that CD74 expressed by activated brain endothelium may cross-present *Pb*A antigens to prime infiltrating CD8^+^ T cells. Accordingly, we assessed the ability of *Pb*A antigen-pulsed, brain-derived endothelium to activate T cells by employing the LR-BSL8.4a reporter T cell line that expresses LacZ in response to the *Pb*A-GAP50 epitope [6]. *Cd74*^−/−^ brain-derived endothelial cells were less able to activate LR-BSL8.4a T cells in the presence of *Pb*A antigens when compared to WT brain-derived endothelial cells (**Figure 2A**). We confirmed these results by isolating brain microvessels from *Pb*A-infected WT and *Cd74*^−/−^ mice at the time of ECM development and incubating them with LR-BSL8.4a reporter T cells for measurement of LacZ expression. Microvessels from *Pb*A-infected WT mice showed a greater ability to cross-present *Pb*A antigens than microvessels from *Cd74*^−/−^ mice (**Figure 2B**). We next assessed cross-presentation of *Pb*A antigens *ex vivo* by using brain microvessels from *Cd8*^−/−^ or *Cd8*^−/−^*Cd74*^−/−^ mice infected with *Pb*A after adoptive transfer with CD8^+^ T cells from WT or *Cd74*^−/−^ mice. The microvessels from recipient *Cd8*^−/−^ mice receiving WT or *Cd74*^−/−^ CD8^+^ T cells from *Pb*A infected mice had a greater ability to cross-present *Pb*A antigens (**Figure 2C**). Together, these results indicate that CD74 expression by brain endothelial cells contributes to cross-presentation of *Pb*A antigens to CD8^+^ T cells and to ECM development.

**Figure 2.**
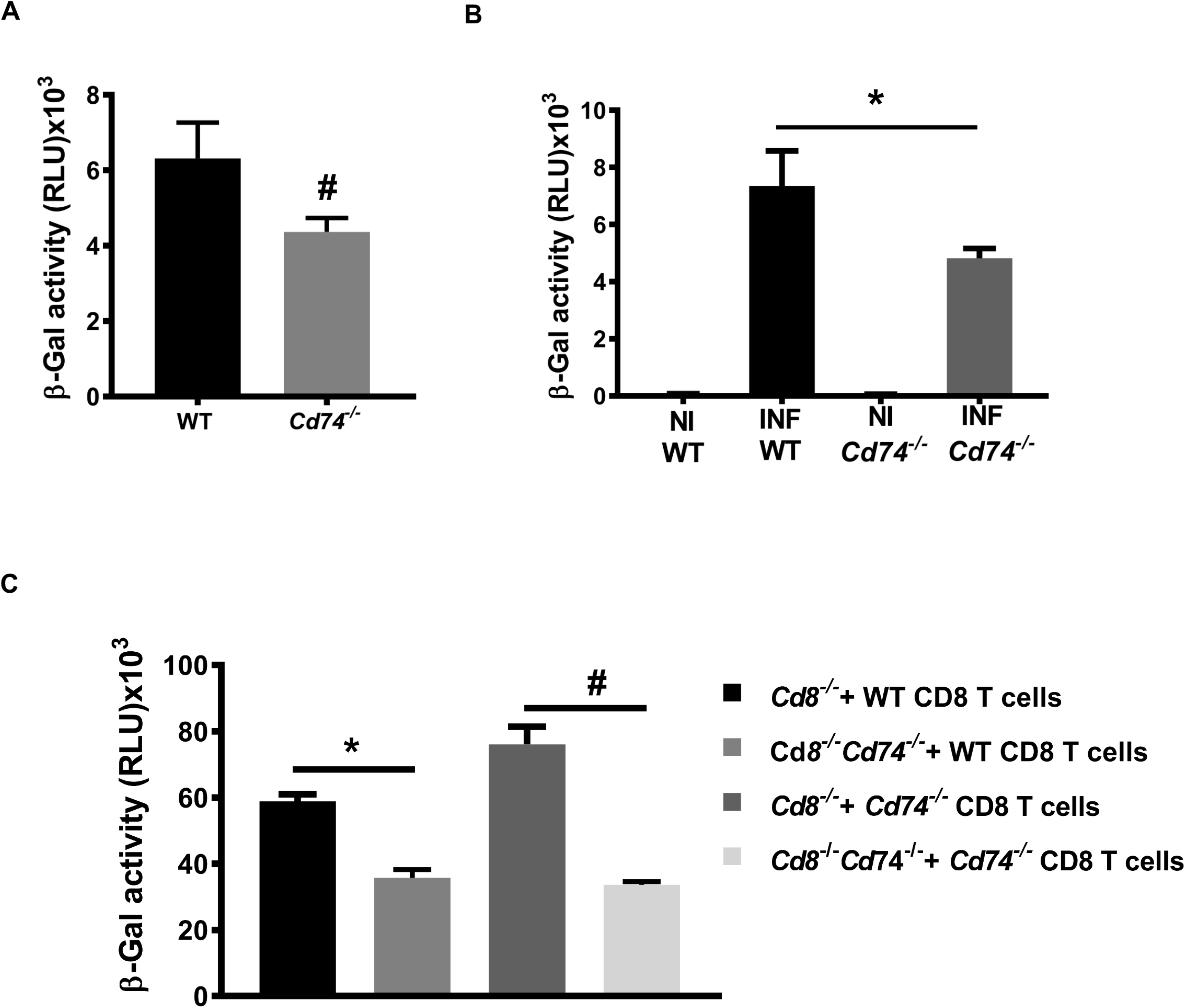
Cross-presentation of *Plasmodium* antigen by brain endothelium is Cd74 dependent. Cross-presentation of *Pb*AGAP_50_ by brain endothelial cells. **A**, BEC isolated from WT and *Cd74*^−/−^ mice were stimulated with 10 ng/ml IFNγ for 24 h, and then incubated for additional 24h with *Pb*A mature stage iRBCS before co-culture with LR-BSL8.4a reporter cells overnight prior to β-galactosidase activity assessment. Data are shown as mean ± SD of three independent biological replicates performed in triplicate; #p<0.0001 by Mann-Whitney test. **B**, Brain microvessel cross-presentation of *Pb*AGAP_50_ from naïve and *Pb*A infected WT and C*d74*^−/−^ mice. Mice were infected with 10^6^ *Pb*A, iRBC and brain microvessels were isolated when WT-infected mice exhibited neurological signs at 7 days after infection and co-incubated with LR-BSL8.4a reporter cells for 24 h and then assessed for β-galactosidase activity. Data are shown as mean ± SD of n=6 mice per group and pooled from two independent biological replicates; **p=0.0021 by Mann-Whitney test. **C**, Brain microvessel cross-presentation of *Pb*AGAP50 from *Cd8*^−/−^ and *Cd8*^−/−^*Cd74*^−/−^ receiving CD8^+^ T cells isolated from *Pb*A infected WT or *Cd74*^−/−^ mice three days before the infection with 10^6^ *Pb*AWT iRBC. Brain microvessels were isolated when the first infected mice exhibited neurological signs at 7 days after infection and co-incubated with LR-BSL8.4a reporter cells for 24 h and then assessed for β-galactosidase activity. Data are shown as mean ± SD of n=5 mice per group and pooled from two independent biological replicates; #p=0.0017 and **p<0.0001 by Kruskal-Wallis test.

### PMIF contributes to the development of ECM by promoting *Pb*A liver stage development

Precedent studies have shown that *Pb*A parasites genetically deficient in PMIF (*Pb*A*mif*-) develop normally in their mosquito hosts and during blood-stage infection [14]. To examine the contribution of PMIF to the development of ECM, we infected C57BL6/J mice with wild-type *Pb*A (*Pb*AWT) or *Pb*A*mif*- iRBC. There was no difference in ECM manifestations, and all the mice succumbed by day seven after infection (**Figure 3A**). Recent studies have shown that liver-stage *Plasmodium* infection is critical for ECM development [17]. We and others recently demonstrated that PMIF is necessary for efficient *Plasmodium* liver-stage development of the parasite and that its absence impairs blood-stage patency [12, 13]. Accordingly, mice infected with *Pb*A*mif*- sporozoites did not exhibit ECM signs and survived until day 25 when compared with mice infected with *Pb*AWT sporozoites; the later mice exhibited neurological symptoms followed by mortality 8-9 days after infection (**Figure 3B**). We measured the expression of CD74 in the brain of infected mice during ECM and observed an increase in *Cd74* mRNA expression in the brains of *Pb*AWT infected mice when compared with brains of *Pb*A*mif*- infected mice (**Figure 3C**). The expression of the inflammatory molecules IFN-γ, perforin, and granzyme B in the brains of mice infected with *Pb*AWT sporozoites also was higher than in mice infected with *Pb*A*mif*- sporozoites (**Figure 3D**).

**Figure 3.**
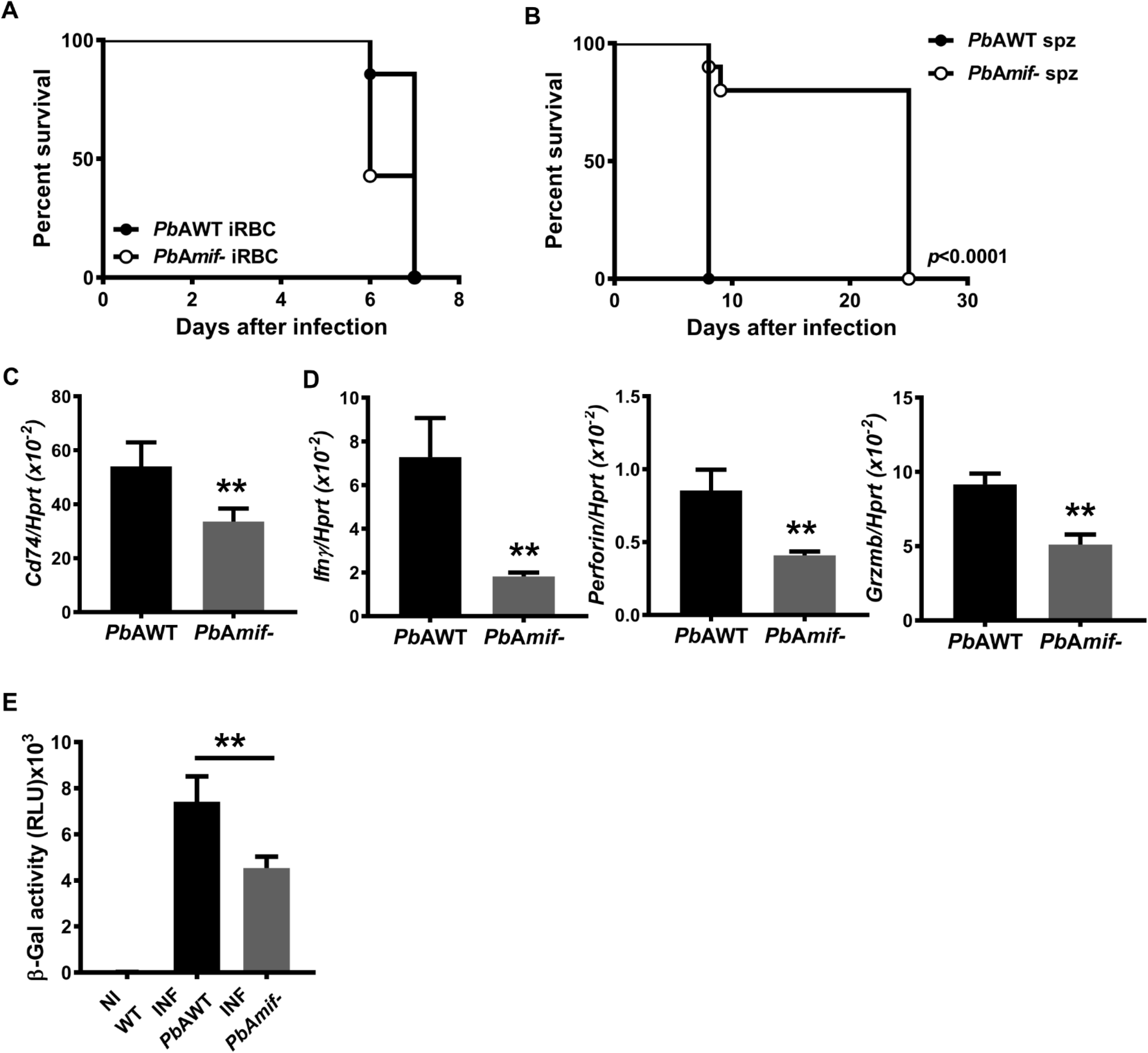
PMIF contributes to the development of ECM. **A**, Kaplan-Meier survival plots for C57BL6/J mice infected with i.p with 10^6^ *Pb*AWT or *Pb*A*mif*- iRBC. Data are form two pooled independent experiments with a total of 6 animals per group. n.s: non-significant by log-rank (Mantel Cox) test. **B**, Kaplan–Meier survival plots for C57BL6/J mice infected with 2×10^3^ *Pb*AWT or *Pb*A*mif*- sporozoites. Data are from three pooled independent experiments with a total of 6 animals per group; p< 0.0001 by log-rank (Mantel Cox) test. WT C57BL6/J mice were infected i.v with 2×10^3^ *Pb*AWT or *Pb*A*mif*- sporozoites. **C,** CD74 transcript expression in the brain of infected mice was measured by q-PCR at day 8 after infection. Results are expressed as mean ± SD of n=6 mice per group pooled from two experiments; **p=0.0022 by Mann-Whitney test. **D**, Transcriptional expression of IFNγ, Perforin and Granzyme B was measured in brain tissue of *Pb*AWT or *Pb*A*mif*- infected WT mice on day 8 after infection by quantitative real-time PCR. Results are expressed as mean ± SD of n=6 mice per group pooled from two experiments; **p=0.0022 for Perforin and IFNγ and **p=0.0317 for Granzyme B, by two-tailed Mann-Whitney test. **E**, Brain microvessel cross-presentation of *Pb*AGAP50 from naïve (NI) and *Pb*AWT or *Pb*A*mif*^−^ infected WT mice. Mice were infected with *Pb*AWT or *Pb*A*mif*-sporozoites and brain microvessels were isolated when *Pb*AWT-infected mice exhibited neurological signs at 7 days after infection. Microvessels then were co-incubated with LR-BSL8.4a reporter cells for 24 h and then assessed for β-galactosidase activity. Data are shown as mean ± SD of n=6 mice per group and pooled from two independent biological replicates; **p=0.0021 by Mann-Whitney test.

To evaluate the contribution of the PMIF/CD74 interaction to *Plasmodium* antigen cross-presentation and ECM development, we infected WT mice with *Pb*AWT or *Pb*A*mif*- sporozoites isolated the brain microvessels at the time of ECM onset, and we incubated them with LR-BSL8.4a reporter T cells. We observed that microvessels from *Pb*AWT infected mice exhibited a greater ability to cross-present *Pb*A antigens than microvessels from *Pb*A*mif*- infected mice (**Figure 3E**). Together, these data confirm the role of PMIF in liver-stage *Plasmodium* infection and the subsequent promotion of inflammation during blood-stage infection and ECM development.

### PMIF promotes *Plasmodium*-infected hepatocyte survival and p53 inhibition through host CD74

Prior studies of *Plasmodium*-infected liver cells have implicated pro-survival roles for hepatocyte growth factor signaling [25, 26] and inhibition of the tumor suppressor p53, which is activated by cellular stress to initiate programmed cell death [27]. Mammalian and parasite MIF molecules promote monocyte survival by increasing p53 phosphorylation at Ser^15^ [9, 11]. Our results suggest that PMIF regulates *Pb*A liver-stage development and promotes the development of ECM. We observed that intracellular parasite content was reduced in HepG2 cells infected with *Pb*A*mif*- sporozoites when compared with *Pb*AWT sporozoites (**Figure 4A**). We used circumsporozoite (CSP) and merozoite surface protein-1 (MSP-1) as indicators of parasite maturation [28]. While CSP was expressed in similar levels **(Figure S2A)**, there was reduced expression of MSP-1 in the HepG2 cells infected with *Pb*A*mif*- parasites, suggesting that PMIF is not necessary for hepatocyte infection but may have a permissive role in pre-erythrocytic parasite development (**Fig S2B**). To assess the mechanistic role of PMIF in liver-stage parasite development, we examined its effect on the survival of infected liver cells by treating infected HepG2 cells with the nitric oxide (NO) donor sodium nitroprusside (SNP) to induce p53 accumulation and apoptosis. We found that HepG2 cells infected with *Pb*A*mif*- sporozoites were significantly more susceptible to NO-induced apoptosis than cells cultured with *Pb*AWT parasites, despite a reduced infection level compared with *Pb*AWT sporozoites (**Figure 4B**). The protection from apoptosis observed in *Pb*AWT infected HepG2 cells was associated with decreased phospho-p53^Ser15^ and intracellular p53 content compared with *Pb*A*mif*- infected cells (**Figure 4C)**. Induction of apoptosis in *Pb*A*mif*- versus *Pb*AWT infected cells also was associated with increased Akt phosphorylation (**Figure S2C**). We confirmed these *in vitro* findings by infecting mice with *Pb*AWT or *Pb*A*mif*- sporozoites. The livers of *Pb*A*mif*- infected mice showed an 80% reduction in parasite burden compared with the livers of *Pb*AWT infected mice, and this was associated with an attendant decrease in expression of the host pro-survival gene *Bcl-2* and an increase in the expression of the pro-apoptotic gene *Bad* (**Figure S2D**).

**Figure 4.**
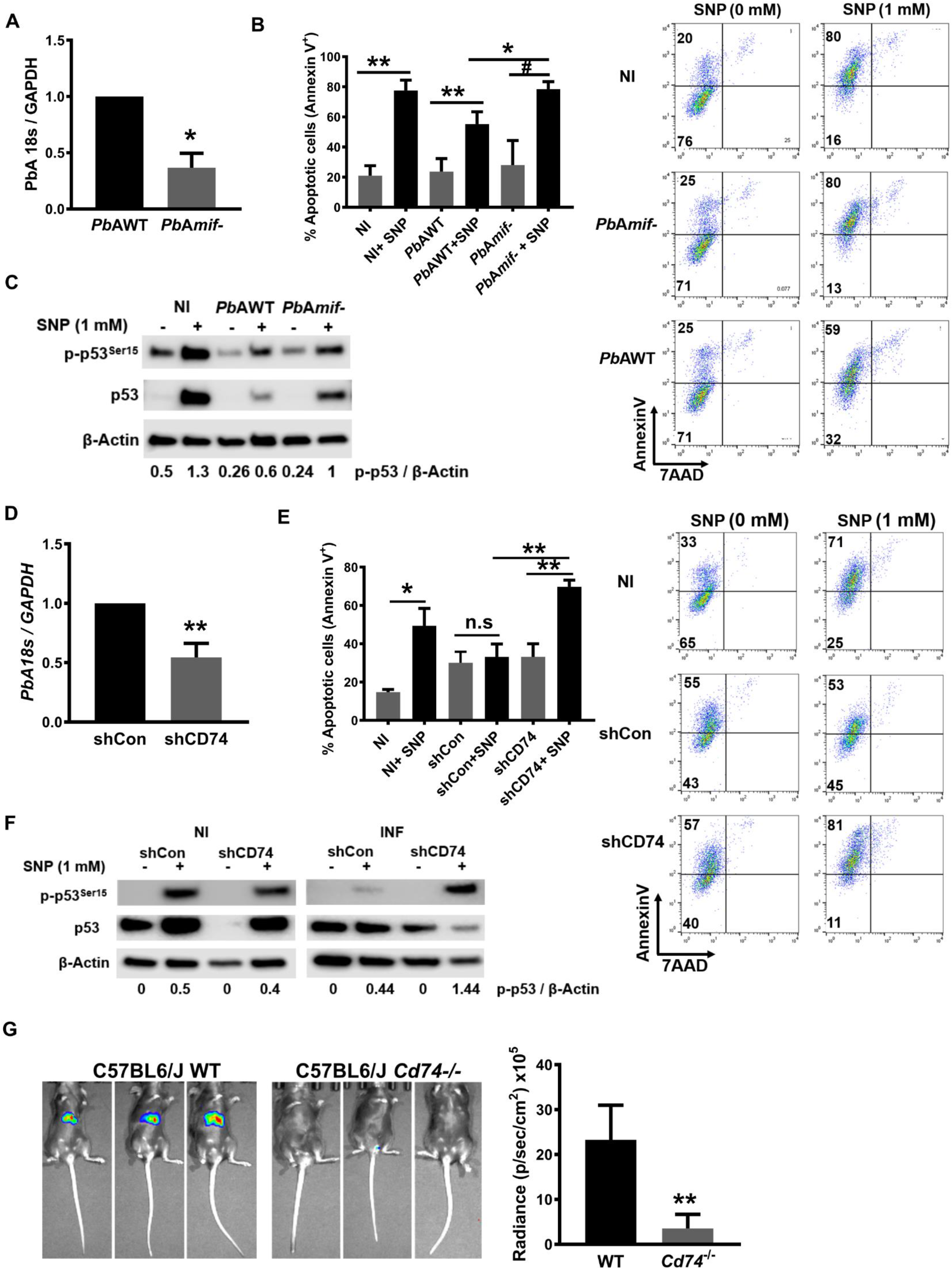
PMIF promotes *Plasmodium*- infected hepatocyte survival and p53 inhibition through host CD74. 1×10^5^ HepG2 cells were infected with 2×10^3^ *Pb*AWT or *Pb*A*mif*- sporozoites. **A**, Parasite load was measured by quantitative PCR of *Pb*A 18S rRNA relative to host GAPDH 48 h after infection. Data are from three independent experiments performed in duplicate. Bars represent the mean ± SD; *p=0.0286 by Mann–Whitney test. 1×10^5^ *Pb*AWT or *Pb*A*mif*- infected HepG2 cells, were cultured for 48 h and treated with 1 mM of the NO donor sodium nitroprusside (SNP) for 4 h to induce apoptosis. **B,** Percentage of apoptotic cells measured by AnnexinV and 7AAD (7-amino-actinomycin D) staining. Data are from three independent experiments performed in duplicate. Bars represent the mean ± SD; *p=0.0133; **p=0.0011; ^Ѱ^p<0.0001; by Kruskal-Wallis test. **C,** Lysates of the same HepG2 cells in C were assessed for total p53 and p53^Ser15^ by Western blotting. NI: non-infected. Numerals represent the mean densitometric scanning ratios. Data are representative of two independent replicates experiments for the western blot analysis. 1×10^5^ HepG2 cells were treated with 10 nM of shRNA directed at CD74 (shCD74) or a control shRNA (shCon), and infected 24 h later with 2×10^3^ *Pb*AWT sporozoites. **D**, Parasite load was measured by quantitative PCR of *Pb*A 18S rRNA relative to host GAPDH. Data are from three independent experiments performed in duplicate. Bars represent the mean ± SD; **p=0.002 by Mann–Whitney test. 1×10^5^ shCon or shCd74 treated HepG2 cells, were infected with 2×10^3^ *Pb*AWT sporozoites and cultured for 48 h. shRNA cells were then treated with 1 mM of the NO donor SNP for 4 h to induce apoptosis **E,** Percentage of apoptotic (measured by AnnexinV and 7AAD staining) *Pb*AWT infected HepG2 cells treated with shCon or shCD74. Data are from three independent experiments performed in duplicate. Bars represent the mean ± SD; **p=0.0095 and n.s.=non-significant; by Kruskal-Wallis test. **F,** Lysates of the same hepatocytes as in E were assessed for total p53 and p53^Ser15^ by Western blotting with β-actin as loading control. NI: non-infected, INF: *Pb*AWT infected. Numerals represent the mean densitometric scanning ratios. Data are representative of two independent replicate experiments for the western blot analysis. **G**, Wild type (WT) or *Cd74*^−/−^ C57BL/6J mice were infected i.v. with 2×10^3^ *Pb*A-luciferase sporozoites and liver *Pb*A-luc load quantified by luminescence at 48 h after infection. Bars represent the mean ± SD; **p=0.0022; by Mann-Whitney test.

We next confirmed the direct role of PMIF signaling through the host MIF receptor by studying sporozoite infection in HepG2 cells after knockdown of CD74 (**Figure S2E**). HepG2 cells treated with shCD74 to reduce CD74 expression had decreased parasite burden compared with treatment with a non-relevant shRNA (shCon) (**Figure 4D**). As expected, *Pb*AWT infected shCD74-treated cells were more susceptible to apoptosis than infected shCon-treated cells (**Figure 4E**). Apoptosis induction also was associated with increased cellular p53^Ser15^ and p53 accumulation in the infected HepG2 cells with reduced CD74 expression (**Figure 4F**). Infection of mice genetically deficient in CD74 (*Cd74*^−/−^) with *Pb*AWT sporozoites revealed a significant reduction in liver burden of *Plasmodium* parasites when compared to WT (*Cd74*^+/+^) mice (**Figure 4G**), and this reduction was associated with a delay in blood-stage patency from 2 to 6 days post-infection (**Figure S2F**). These results support the essential role of CD74 in mediating PMIF action and promoting *Plasmodium* pre-erythrocytic development leading to blood-stage infection.

### Pharmacologic PMIF antagonism reduces *Pb*A infection and protects against cerebral malaria

Our experimental results support a central role in malaria infection for host CD74 and its activation by PMIF to promote the survival of infected hepatocytes, leading to inflammatory blood-stage infection and subsequent ECM pathophysiology. Anti-PMIF antibodies have been reported in malaria patients [29] and we assessed the ability of malaria infected sera to interfere with PMIF binding to CD74 using an established ELISA employing the recombinant CD74 ectodomain [19, 30]. Such sera inhibited PMIF binding to CD74 when compared to sera from uninfected healthy controls (**Figure 5A**). Moreover, sera from patients with clinically uncomplicated malaria were more effective in reducing PMIF/CD74 interaction than sera from those with complicated malaria (*e.g.,* severe anemia, cerebral malaria [15]), suggesting that a more effective anti-PMIF serologic response may be associated with reduced inflammatory sequelae during infection.

**Figure 5.**
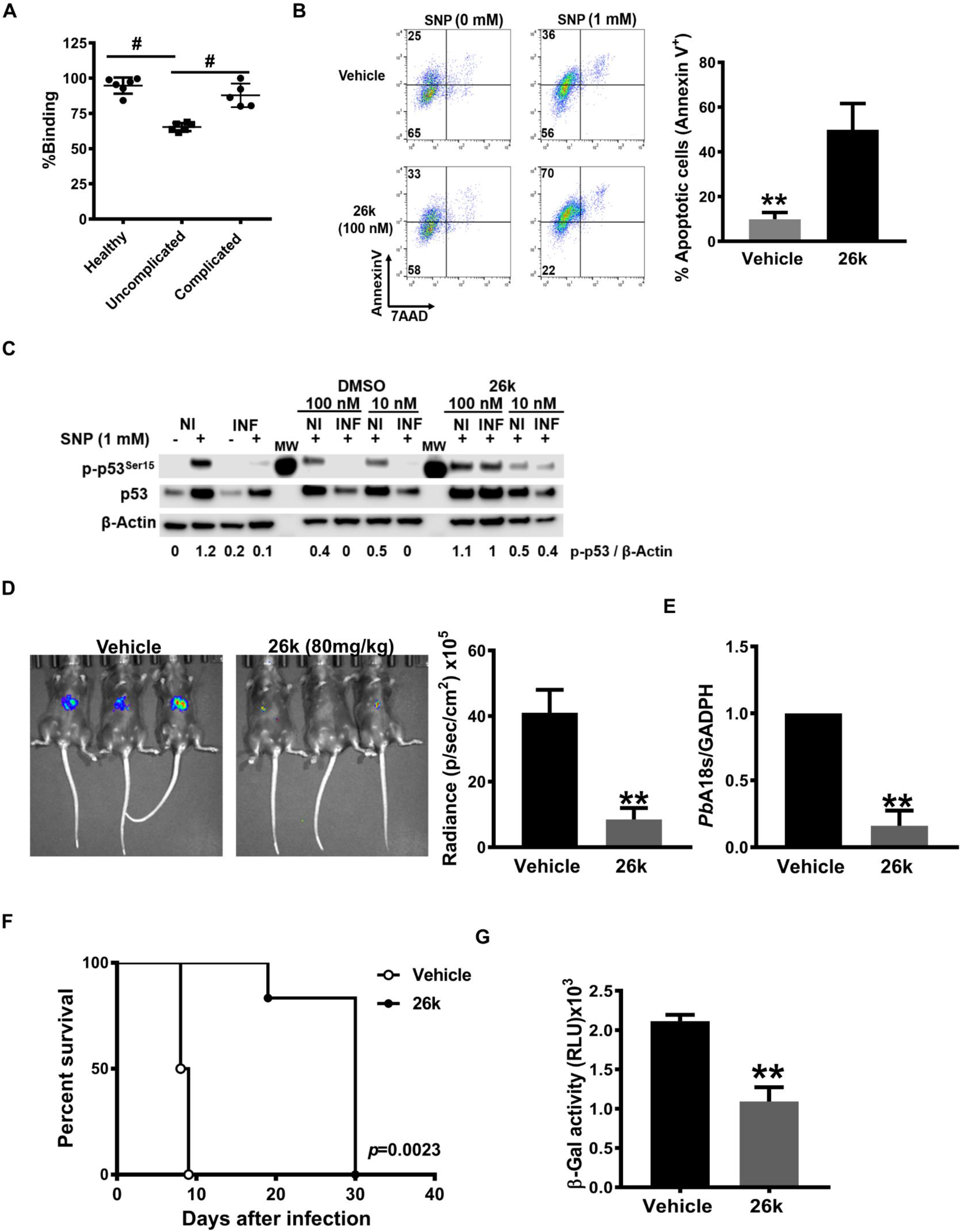
Inhibition of PMIF/CD74 axis is associated with protection from severe malaria. **A.** Effect of human serum on PMIF binding to the immobilized human CD74 ectodomain (aa 134-232). Measured values are relative to control without serum for each condition (n=6 healthy uninfected controls, n=6 uncomplicated malaria, n=6 complicated malaria. Mean±SD; #p<0.0001 by 1-way ANOVA. **B,** Percentage of apoptotic *Pb*AWT infected HepG2 cells measured by AnnexinV and 7AAD staining after 26k or vehicle treatment. 1×10^5^ *Pb*AWT infected hepatocytes treated with 26K (100 nM) or vehicle were cultured for 48 h followed by the addition of the NO donor SNP (1 mM) for 4 h to induce apoptosis. Data are from three independent experiments performed in duplicate. Bars represent the mean ± SD; *p=0.0011; by two-tailed Mann-Whitney test. **C,***Pb*AWT infected HepG2 cells were lysed and assessed for total p53 and p53^Ser15^ by Western blotting with β-actin as loading control. Numerals represent the mean densitometric scanning ratios. Data are representative of two independent replicate experiments for the western blot analysis. NI: non-infected, INF: *Pb*A infected. C57BL/6J mice were treated with vehicle or 26k (80 mg/kg, ip) before (0 h), 24 h, and 48 h after i.v. infection with 2×10^3^ *Pb*A-luciferase sporozoites. **D,** Liver *Pb*A-luc burden was quantified by luminescence and **E,** by qPCR of liver *Pb*A 18S rRNA relative to host GAPDH 48 h after infection. Bars represent the mean ± SD; **p=0.0043 (**D**), **p=0.0079 (**E**); by Mann-Whitney test. **F**, Kaplan–Meier survival plots for vehicle and 26k treated mice following infection with *Pb*A-luc sporozoites. Data are from two pooled independent experiments with 6 animals per group; **p = 0.0023 by log-rank (Mantel Cox) test. **G**, Brain microvessel cross-presentation of *Pb*AGAP50 from vehicle and 26k treated mice following infection with PbA-luc sporozoites. Brains microvessels were isolated when vehicle-treated, infected mice exhibited neurological signs at 9 days after infection and co-incubated with LR-BSL8.4a reporter cells for 24 h and then assessed for β-galactosidase activity. Data are shown as mean ± SD of n=6 mice per group and pooled from two independent biological replicates; **p=0.0022 by two-tailed Mann-Whitney test.

Small molecule MIF inhibitors have been developed and are in clinical evaluation [31, 32]. We recently identified a small molecule PMIF antagonist, termed 26k, that shows a 2500-fold greater selectivity for PMIF than for host MIF (*K_i_* = 40 nM for PMIF versus Ki >100 µM for MIF) [18] [19] and blocks downstream ERK1/2 MAPK signaling (**Figure S3A**). We treated *Pb*A sporozoite infected HepG2 cells with 26k *in vitro* and measured parasite content by expression of *Pb*A18s RNA together with sensitivity to apoptosis induction. Parasite burden decreased in the cells treated with 26k compared with vehicle (**Figure S3B**). Hepatocytes infected with *Pb*A and treated with 26k also were more susceptible to apoptosis, as evidenced by Annexin V staining (**Figure 5B**). Moreover, 26k-treated cells showed increased p53 phosphorylation and intracellular accumulation, a known consequence of CD74 blockade (**Figure 5C**) [9].

To determine the potential *in vivo* action of 26k, we treated C57BL/6J mice with 26k before infection with 2×10^3^ *Pb*A sporozoites and then continued treatment once daily for 2 days. We assessed liver-stage infection, blood-stage patency, and the development of ECM and lethality. Treatment with 26k markedly decreased parasite burden in the liver at 48 h after infection when compared with vehicle controls (**Figure 5D and E**). Sporozoite infection led to blood-stage patency after 3 days in vehicle-treated mice but not until day 5 in the 26k treated group (**Figure S3C**). All vehicle-treated mice developed ECM symptoms (head deviation, ataxia, and paraplegia) 8 days after sporozoite infection, and all mice succumbed to cerebral malaria by days 9-10. By contrast, all mice treated with 26k were spared from cerebral malaria symptoms and did not succumb until after day 20 (**Figure 5F**, and S3D). We also examined the impact of 26k on *Plasmodium* antigen cross-presentation by brain microvascular endothelial cells. Brain microvessels from *Pb*A infected WT mice were isolated at the time of ECM and treated *in vitro* with 26k (or vehicle) together with LR-BSL8.4a reporter T cells. Microvessels treated with 26k showed a reduction in *Pb*A antigen cross-presentation as quantified by LacZ expression (**Figure 5G**). We also examined if 26k administration *in vivo* prevented ECM when mice were inoculated directly with iRBCs. We observed only partial protection from ECM in mice treated with 26k in this model (**Figure S3E**).

Taken together, these results support the conclusion that pharmacologic inhibition of the PMIF/CD74 interaction may be a promising approach to protect from liver infection and, consequently, ECM.

## Discussion

*Plasmodium* parasites have evolved highly specialized invasion strategies that function to evade immune destruction, sustain infection, and ensure completion of their life cycle. Our study highlights the importance of the pre-erythrocytic phase of infection in the development of the immune response and the subsequent progression of ECM. The present findings also emphasize the importance of investigating *Plasmodium* genes whose pathologic relevance may be underestimated based on the stage of the infection.

Our precedent studies reported that PMIF promotes inflammatory signaling through the host MIF receptor CD74 [15]. We show herein that PMIF increases the expression of CD74 in the brain of infected mice and serves a previously unforeseen role in the cross-presentation of *Plasmodium* antigens to promote a CD8+ T cell-mediated, pathologic inflammatory response. Increased IFNγ expression also is a feature of ECM pathology [33] and may further contribute to brain endothelial CD74 expression [34]. While dendritic cells are considered the primary antigen-presenting cells responsible for activating CD4 and CD8 T cell responses against *Plasmodium* [35], endothelial cell activation contributes importantly to blood-brain barrier breakdown and neurological disease [6, 22]. Our data further implicate PMIF, which appears to be universally expressed by the *Plasmodium* genus [30, 31], and its interaction with hepatocyte CD74 as an adaptive mechanism for sporozoites to usurp a host-protective apoptosis pathway to prevent their destruction and enable differentiation and patent infection.

Independent genetic studies indicate that relative increases in liver CD74 expression correlate with enhanced susceptibility to hepatocyte infection by sporozoites [16]. Immunostaining studies suggest that PMIF is expressed on the surface of infective sporozoites and within the parasitophorous vacuole during liver stage development [12, 14]. The precise localization of PMIF interaction with host CD74 is currently unclear; however, we suggest two possible scenarios: 1) by contact between PMIF on the invading sporozoite and CD74 expressed on the hepatocyte cell surface, or 2) after sporozoite internalization and contact between PMIF in the parasitophorous vacuole and endosomal-expressed CD74 during later-stage *Plasmodium* development [36]. Either pathway could initiate activation of Akt and cellular pro-survival pathways [27]. It is also notable that PMIF vaccination is associated with a more robust liver resident memory CD8+ T cell response, suggesting that the apoptotic destruction of infected hepatocytes promotes the development of protective immunity [13].

We additionally show that the PMIF/CD74 interaction pathway is amenable to pharmacologic targeting. The PMIF selective antagonist 26k [28,29] recapitulates the experimental effects of genetic PMIF or CD74 deficiency, and short term administration of 26k reduced *Pb*A intrahepatic development and provided complete protection against cerebral malaria. Together with precedent co-crystallization studies supporting the selectivity of 26k for PMIF versus host MIF, these results provide proof-of-concept for pharmacologic PMIF antagonism as a tractable approach for malaria prophylaxis or liver-stage treatment, and potentially across a range of *Plasmodium* species and strains [29]. PMIF is highly conserved among *Plasmodium* species, with only five single nucleotide polymorphisms in PMIF among the 202 sequenced strains of *P. falciparum* present in the PlasmoDB resource [9, 10]. This high degree of structural conservation may be auspicious for therapeutic targeting, particularly in a genetically complex pathogen prone to resistance development. Additional studies to optimize the absorption, distribution, metabolism, and excretion properties of 26k will be necessary to advance PMIF selective inhibitors such as 26K into clinical utility.

## Materials and Methods

### Mice

Female WT or *Cd74*^−/−^ C57BL/6J mice between 6-10 weeks of age were purchased from The Jackson Laboratory and used for the study. *Cd8*^−/−^*Cd74*^−/−^ mice were obtained by crossing *Cd8*^−/−^ with *Cd74*^−/−^ mice. Swiss Webster mice were obtained from The Jackson Laboratory. All animals were maintained in a specific pathogen-free facility at Yale Animal Resource Center. All animal procedures followed federal guidelines and were approved by the Yale University Animal Care and Use Committee, approval number 2017-10929.

### Parasites and infection

*Pb*AWT (MR4), *Pb*A*mif*- (Leiden Malaria Group) [14], or *Pb*AWT-GFP-luciferase (MR4) parasites were cycled between Swiss Webster mice and *Anopheles stephensi* mosquitoes. For erythrocytic infection, cryopreserved stocks of infected red blood cells (iRBCs) were injected (10^6^ iRBCs/mouse), and blood parasitemia was monitored by Giemsa-stained blood smears and flow cytometry [13]. For the pre-erythrocytic stage infection, salivary gland sporozoites were extracted from infected mosquitoes on day 19 post-blood meal infection. WT or *Cd74*^−/−^ C57BL/6J mice were infected by i.v. tail injection of 2000 *Pb*AWT, *Pb*A*mif*- or *Pb*AWT-GFP-luciferase sporozoites, and blood patency was monitored beginning day 3 by blood smear and flow cytometry. Liver parasite burden was monitored at 48 h after infection using an IVIS imaging system (Caliper) or quantitative PCR [13].

For adoptive transfer, splenocytes were isolated six days after infection of WT or *Cd74*−/− mice infected with 10^6^ *Pb*AWT iRBC, CD8 T cells were purified with anti-CD8 (Ly-2, Miltenyi Biotech) according to the manufacturer’s protocol. 1×10^7^ cells were transferred, i.v. into recipient C57BL6/J *Cd*8^−/−^ or *Cd*8^−/−^*Cd74*^−/−^ mice and mice infected three days after with 10^6^ *Pb*AWT iRBC.

### Hepatocyte infection, apoptosis induction, Annexin V assay, and quantification of p53 by western blotting

For apoptosis assessment, 1×10^5^ HepG2 cells (ATCC) were seeded in complete EMEM medium (ATCC) (10% FBS (Atlanta Biologicals), 1% streptomycin/penicillin (Thermo-Fisher), and infected with 2 ×10^3^ *P*bAWT or *Pb*A*mif*- sporozoites. 48 h after infection, cells were treated with 1 mM of SNP (Sodium Nitroprusside, Sigma) for 4 h or left untreated as a control. For PMIF pharmacologic inhibition, cells were treated with 26k (10 nM, 100 nM) or with an equivalent concentration of vehicle (0.1% DMSO) (Sigma) before infection. Cells then were detached with Acutase (MP Bio), and cell suspensions split for Western blot or Annexin V analysis. For Annexin V analyses, cells were stained with Pacific Blue-Annexin V and 7AAD (7-aminoactinomycin D) (Biolegend) before running in an LSRII cytometer (BD Biosciences). For quantification of p53 by western blotting, *Pb*AWT-infected HepG2 hepatocytes were detached with Accutase (MP Biolabs) and pelleted. Western blots were performed by lysing cell pellets in RIPA buffer (ThermoFisher) according to standard protocols and using antibodies directed against p53-Ser^15^ (clone 1C12) or total p53 (pAb) (Cell Signaling Technology). For quantification, density signals were normalized to an anti-β-actin Ab (LICOR) and developed with anti-mouse or anti-rabbit Abs conjugated with HRP (LI-COR Biosciences). Membranes were visualized using an Odyssey-Fc imaging system (LI-COR Biosciences). Each western blot panel was developed from the same membrane that was re-probed after stripping.

### Quantification of liver-stage *Pb*AWT and *Pb*Amif- infection, and *Cd74* knockdown

HepG2 liver cells infected with *Pb*AWT or *Pb*A*mif*- sporozoites were lysed at 24 h or 48 h after infection, cellular proteins transferred to PVDF membranes (Millipore), and analyzed by western blotting using anti-CSP (MRA-100) and anti-MSP-1 (MRA-667) antibodies obtained from MR4 ATCC (Manassas, VA). β-actin was used as a loading control. For treatment with siRNA, hepatocytes were transfected with 15 pmol of siRNA (Ambion) targeting CD74 mRNA (3 target sequences in exon 2) or siCtrl (scrambled unrelated sequence) as a negative control. siRNA was complexed with Lipofectamine RNAimax reagent (ThermoFisher) and added to the cells for 24 h; cells then were infected with *Pb*AWT sporozoites.

### Murine *Pb*A infection and 26k treatment

C57BL/6J mice received three i.p. injections of 26k (4-(3-methoxy-5-methylphenoxy)-2-(4-methoxyphenyl)-6-methylpyridine) [18], synthesized by CheminPharma LLC (Branford, Connecticut), at 80 mg/kg after dissolution in PEG400 (Sigma-Aldrich) in a sonicating water bath. HP-P-cyclodextrin (Sigma-Aldrich) was added to prepare a 4 mg/ml solution. Control mice received vehicle alone. Immediately after the first i.p. injection, mice were infected i.v. with 2×10^3^ *Pb*A-luc-GFP sporozoites; the second and third injections of 26k were given 24 h and 48 h later, and always after measurement of *Pb*A liver burden. The *Pb*A liver burden was quantified 48 h after infection by luminescence emission after luciferin injection (Perkin Elmer) using an IVIS apparatus (Caliper). Livers were excised 48 h after infection, the total RNA extracted and purified with Trizol (Life Technologies), and parasites quantified by RT-qPCR using primers for *Pb*18s.

### Brain microvessel cross-presentation assay

WT, *Cd74^−/−^*, *Cd8*^−/−^ or *Cd8*^−/−^*Cd74*^−/−^ C57BL/6J mice were infected by intravenous injection of sporozoites or by intraperitoneal injection of *Pb*A infected red blood cells. Infected mice were sacrificed when the signs of ECM (head deviation, ataxia) were manifested. Control naïve mice were sacrificed contemporaneously with the experimental group. The technique for isolating brain microvessels and quantification of cross-presentation of the parasite-derived GAP50 epitope by LR-BSL8.4a reporter T cells [22] was performed according to the published protocol [21]. Quantification of β-galactosidase activity by activated LR-BSL8.4a reporter cells was performed by using a luminescence β-galactosidase assay (ThermoFisher) and the resultant signal quantified with a microplate reader (Tecan).

### Isolation of Brain-infiltrating lymphocytes for flow cytometry

Naïve or infected WT and C*d74*^−/−^ C57BL/6J mice were sacrificed and perfused intra-cardially with 20 ml 1x DPBS (’ ‘Dulbecco’s Phosphate Buffer Saline). Brains were minced in RPMI, digested with 10 µg/ml of DNase I (Sigma) and 0.5 mg/ml for 30 min at 37°C, and homogenized with a pestle then filtered with a 70 µm cell-strainer (BD Falcon). The suspension then was centrifuged at 200xg for 5min, and the pellet re-suspended in 90% Percoll (GE Healthcare) and overlaid with a 70%, 50% and 30% Percoll gradient. After centrifugation at 500xg for 10 min, the cell interphase was collected treated with RBC lysis buffer, washed once, and re-suspended in complete RPMI medium (containing 10% FBS (Atlanta Biologicals) and 1% penicillin-streptomycin (Thermo Fisher)). 100 µl of brain cell suspensions were stimulated with 1 µM of peptide (SQLLNAKYL) in the presence of 10 µg/ml Brefeldin A. After 5 h incubation at 37°C, the cells were centrifuged and washed once with 100 µl DPBS + 5% FBS. The cells then were re-suspended and incubated with FITC-labelled SQLLNAKYL-H-^2b^ tetramer for 15 min on ice before staining with anti-CD8-PE Cy7 (clone 53-6.7, Biolegend), anti-CD4-PerCP Cy 5.5 (clone RM4, Biolegend), and anti-CD11a antibodies in the presence of FCR-block for 20 min on ice. Cells were washed, pelleted, and permeabilized by re-suspension in 100 µl Fix/Perm buffer on ice for 15 min. Cells then were washed once with 1x Perm Wash buffer (BD Bioscience) and stained with anti-IFN-γ-APC-Cy7 (clone XMG1.2, Biolegend) and GrmzB-Pe-Cy7 (clone NGZB, eBioescience) in 1x Perm Wash buffer (BD Bioesciences) for 20 min on ice. Finally, cells were washed, centrifuged once, and re-suspended in 200 µl PBS+5% FBS for flow cytometry. The data were acquired on an LSRII flow cytometer (BD Biosciences) and analyzed using the FlowJo software (version 10).

### Patient Samples and PMIF-CD74 binding assay

Sera from a previously characterized Zambian cohort of *P. falciparum*-infected patients were used in the study [15, 37]. The interaction between PMIF and CD74 was analyzed as previously described [19, 30]. Briefly, 96-well plate were coated with 26 ng/ml of recombinant CD74 ectodomain (aa 114-243) in PBS and incubated overnight at 4°C. After washing with PBS/0.1% Tween-20, the plate was blocked with Superblock reagent (Pierce) for 2 hours. Biotinylated recombinant PMIF (5ng/ml) was incubated for 45 min, with human serum (diluted 1:1000) from a previously described repository of healthy donors or subjects with uncomplicated or complicated malaria [15]. After washing and incubating with Streptavidin-HRP (Roche), the peroxidase substrate is 3,3’-5,5’-tetramethylbenzidine (TMB, Roche) was added, and after 20 min incubation, the reaction was stopped with 1N H_2_SO_4_/HCl. The results were expressed as the percentage of binding in the presence versus the absence of serum.

### Statistical analysis

All statistical analysis was performed as described before (15), using Software Prism v.6.0, (GraphPad). Statistical significance was indicated at *p* values of less than 0.05, 0.01, or 0.001. All data were expressed as a mean ± SD of at least two independent experiments. Mouse survival times were analyzed by the Mantel-Cox log-rank test. All other data were first tested for Gaussian distribution of values using a D’Agostino-Pearson normality test. The statistical significance of differences was assessed using the Kruskal-Wallis or Mann–Whitney *U* test for non-parametric data distribution and ANOVA or Student’s *t*-test for parametric data.

### Ethics Approval

All animal procedures followed federal guidelines and were approved by the Yale University Animal Care and Use Committee, approval number 2017-10929. De-identified human sera collected from a prior study were used for in vitro ELISA studies (Yale HIC #0804003730).

## Supporting information

Uncropped western blot

## Figures Legends

**Figure S1.**
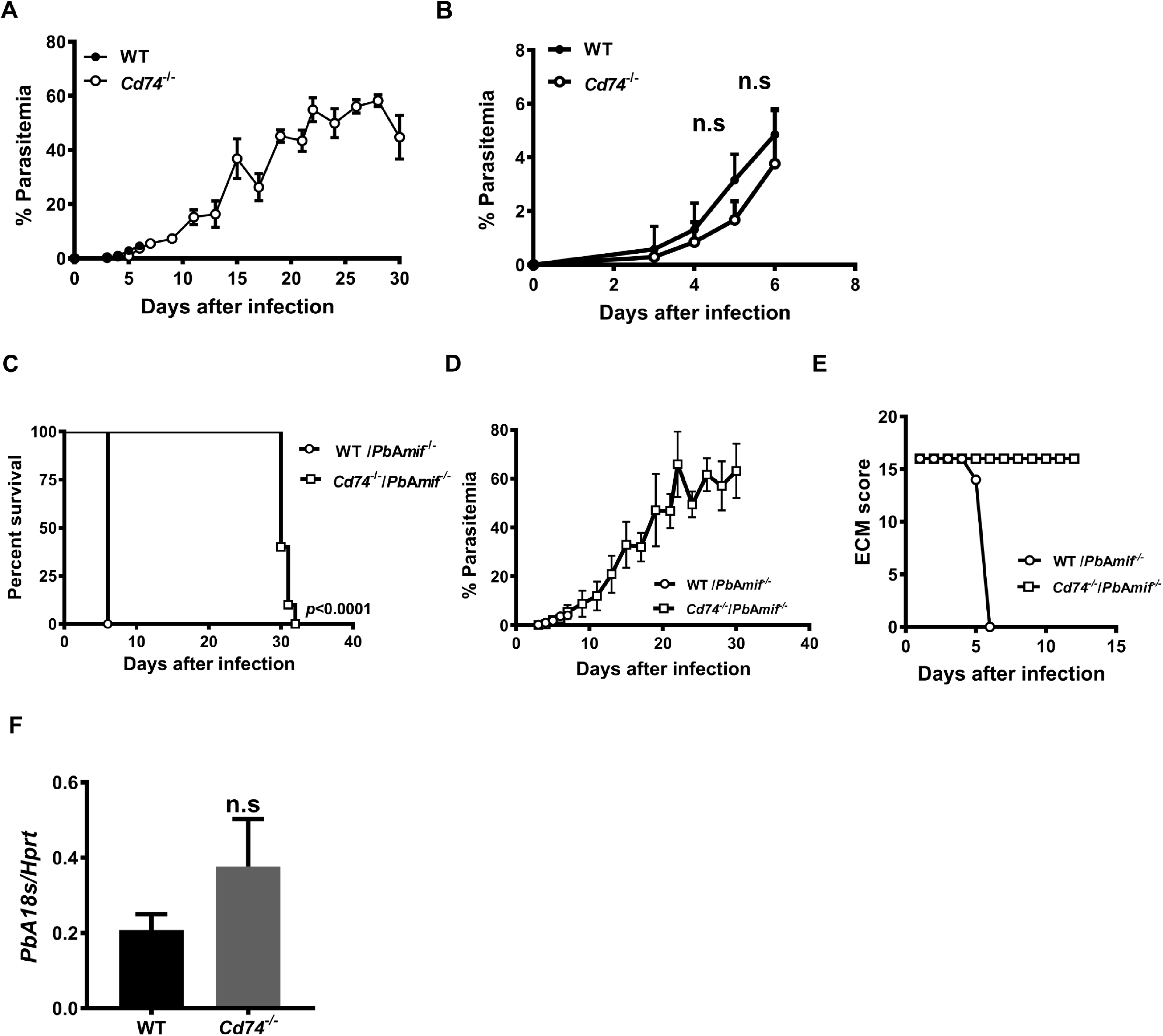
CD74 is necessary for the development of ECM. **A**, Peripheral blood parasitemia for CD74 deficient (*Cd74^−/−^*) and WT C57BL6/J mice infected i.p with 10^6^ *Pb*A iRBC. **B**, Peripheral blood parasitemia during the 6 first days of infection. Data are shown as mean ± SD of n=10 WT and n=10 *Cd74*^−/−^ mice and pooled from three independent experiments. Wild type (WT) and *Cd74*^−/−^ C57BL6/J mice were infected i.p with 10^6^ *Pb*A*mif*^−^ iRBC **C**, Kaplan–Meier survival plots for WT and *Cd74*^−/−^ mice following infection with *Pb*A*mif*^−^. Data are from two pooled independent experiments with 10 animals per group; p < 0.0001 by log-rank (Mantel Cox) test **D**, peripheral blood parasitemia. **E**, ECM malaria score was assessed as described before. Data are shown as mean ± SD of n=7 WT and n=10 *Cd74*^−/−^ mice and pooled from 2 independent experiments. **F**, *Pb*A18s transcript expression in the brain of infected WT and *Cd74*^−/−^ measured by quantitative real-time PCR 7 days after infection was measured in brains tissue of *Pb*A infected WT and *Cd74*^−/−^ mice on day 7 after infection by quantitative real-time PCR. Results are expressed as mean ± SD of n=6 mice per group pooled from two experiments; n.s.: non-significant by Mann-Withney test.

**Figure S2.**
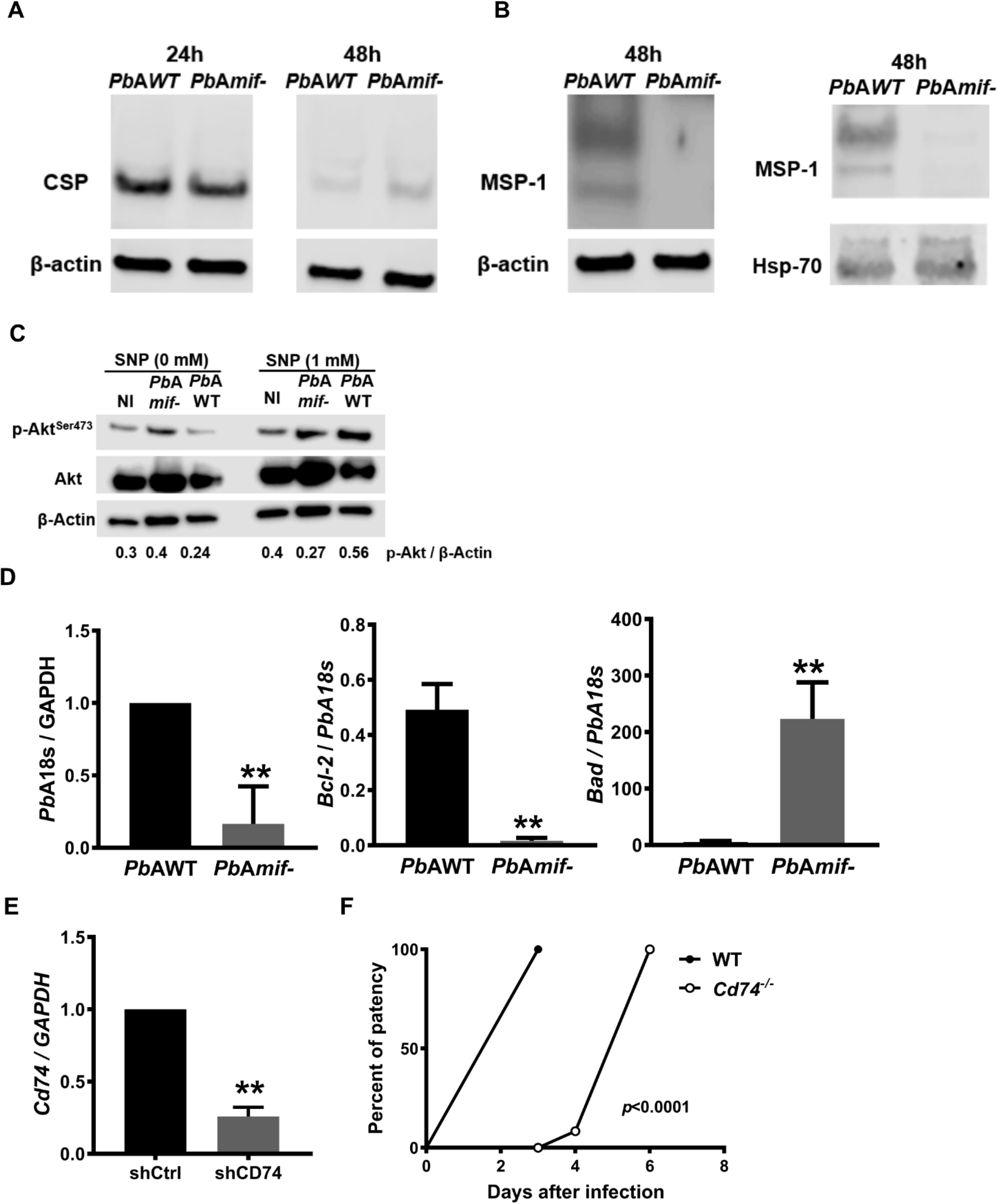
PMIF influences *Pb*A liver-stage development and promotes survival of *Plasmodium*-infected hepatocytes by inhibiting p53 activity. 1×10^5^ HepG2 cells were infected with 2×10^3^ *Pb*AWT or *Pb*A*mif*- sporozoites. Hepatocellular content of A, CSP and **B**, MSP-1 at 24 h and 48 h after infection with 2×10^3^ *Pb*AWT or *Pb*A*mif*- sporozoites assessed by western blot relative to *β*- actin as loading control. Hepatocellular content of MSP-1 at 48 h after infection with 2×103 *Pb*AWT or *Pb*A*mif*- sporozoites was also assessed by western blot relative to *Pb*AHSP70 as loading control. Cultured hepatocytes (1×10^5^ HepG2 cells/well) infected with 2×10^3^ *Pb*AWT or *Pb*A*mif*- sporozoites followed by the addition of 1 mM SNP at 4 h and 48 h after infection. **C**, Lysates were assessed for total Akt and pAkt^Ser473^ by Western blotting with β-actin as loading control. Numerals represent the densitometric scanning ratios. **D**, Transcriptional expression of IFNγ, Perforin and Granzyme B was measured in brain tissue of *Pb*AWT or *Pb*A*mif*- infected WT mice on day 8 after infection by quantitative real-time PCR. Results are expressed as mean ± SD of n=6 mice per group pooled from two experiments; **p=0.0022 for Perforin and IFNγ and **p=0.0317 for Granzyme B, by two-tailed Mann-Whitney test. 1×10^5^ HepG2 cells were treated with 10 nM of shRNA directed at CD74 (shCD74) or a control shRNA (shCon), and infected 24 h later with 2×10^3^ *Pb*AWT sporozoites. **E,***Cd7*4 mRNA expression measured by qPCR in HepG2 after 48h treatment. Data are from three independent experiments performed in duplicate. Bars represent the mean ± SD; **p= 0.002 by Mann-Whitney test. Wild type (WT) or *Cd74*^−/−^ C57BL/6J mice were infected i.v. with 2×10^3^ *Pb*A-luciferase sporozoites and liver *Pb*A-luc load quantified by luminescence at 48 h after infection. Bars represent the mean ± SD; **p=0.0022; by Mann-Whitney test. **F**, Kaplan-Meier plots showing the percentage of WT (●) and *Cd74*−/− (○) C57BL/6J mice with blood-stage patency following i.v. infection with 2×10^3^ *Pb*A-luc sporozoites. Patency was determined by microscopic enumeration of thin blood smears. Data are from two independent experiments, with 3-4 animals per group; p<0.0001 by Log-rank (Mantel Cox) test.

**Figure S3.**
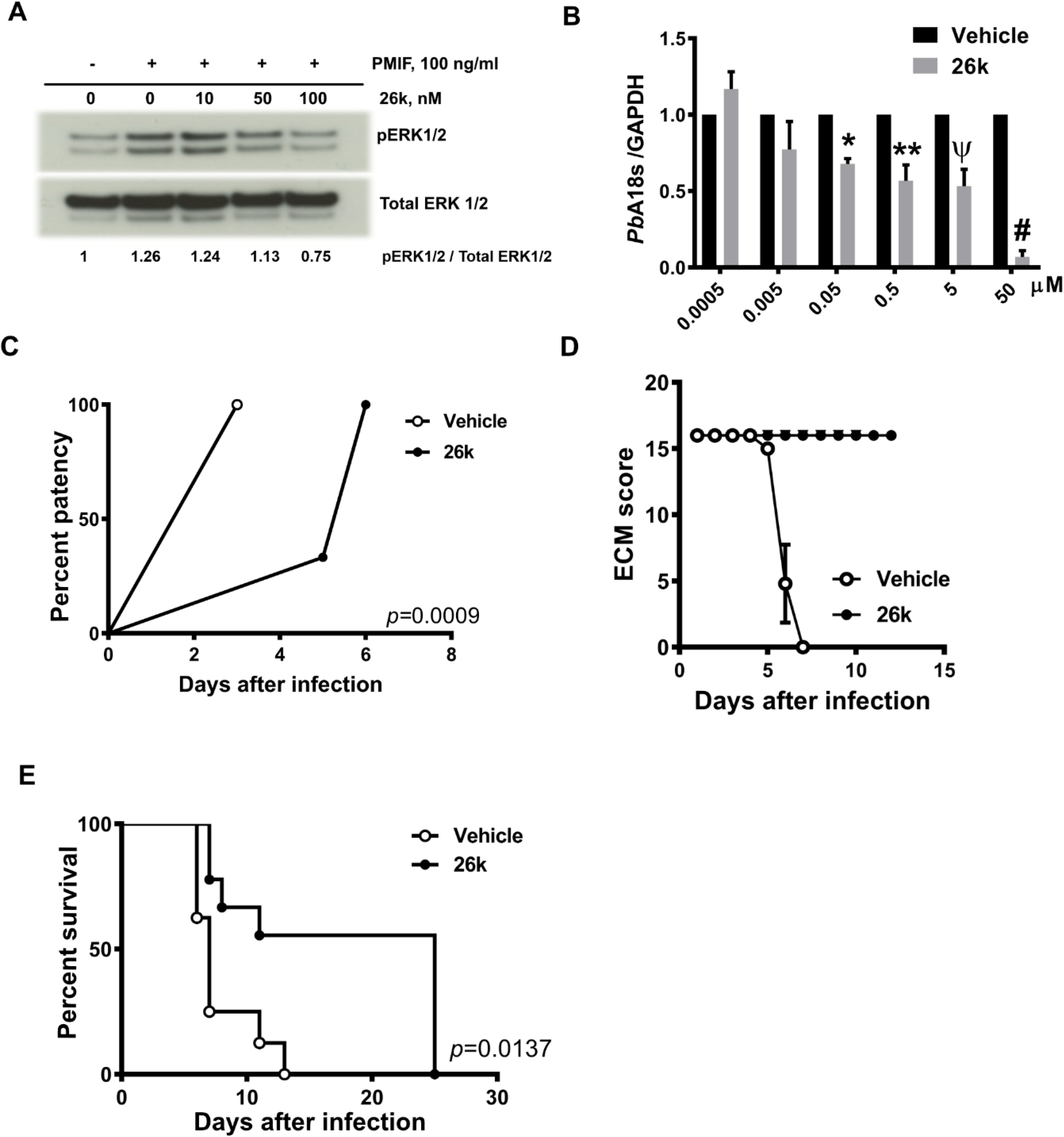
The small molecule PMIF antagonist 26k reduces PMIF/CD74 signal transduction and protects from ECM. **A.** BMDM were treated with or without recombinant PMIF pre-incubated with vehicle control (DMSO) or small molecule 26k (10,50 or 100 nM) for 2h. Cells were lysed and the lysates assessed for total ERK and pERK^Thr202/Tyr204^ by western blotting. Numerals represent the densitometric scanning ratios. HepG2 cells (1×10^5^ cells/well) were infected with 2×10^3^ *Pb*AWT sporozoites and treated with 26k or vehicle. **B**, Hepatocellular parasite load was measured by quantitative PCR of *Pb*A 18S rRNA relative to host GAPDH 48 h after treatment with 26k (0.5 nM to 50 µM) or vehicle. Data are from three independent experiments performed in duplicate. Bars represent the mean ± SD; *p=0.0336, **p=0.0021, Ѱp=0.0008, #p<0.0001 by Kruskal-Wallis test. C57BL/6J mice were treated with vehicle or 26k (80 mg/kg, ip) before (0 h), 24 h, and 48 h after i.v. infection with 2×10^3^ *Pb*A-luciferase sporozoites**. C.** Kaplan-Meier plots showing the percentage of vehicle (o) and 26k (●) treated mice with blood-stage patency (determined by microscopic enumeration of thin blood smears) following i.v. infection with 2×10^3^ *Pb*A-luc sporozoites and **D**, ECM malaria score. Data are from two pooled independent experiments, with 6 animals per group p<0.0001 by Log-rank (Mantel Cox) test. C57BL/6J mice were treated with vehicle or 26k (80 mg/kg, ip) before (0 h) and then every two days after i.p infection with 10^6^ *Pb*AWT iRBCs**. E**, Kaplan– Meier survival plots for vehicle (o) and 26k (●) treated mice following infection with *Pb*A iRBC. Data are from two pooled independent experiments with 8 total animals per group; p = 0.0137 by log-rank (Mantel Cox) test.

