## Supplementary material for "Suppression of *Plasmodium* MIF-CD74 Signaling Protects Against Severe Malaria": Uncropped western blot

#### Uncropped blots for Fig 1D

CSP

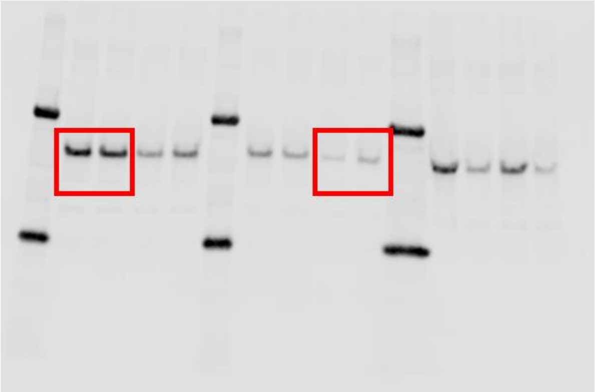

β-Actin

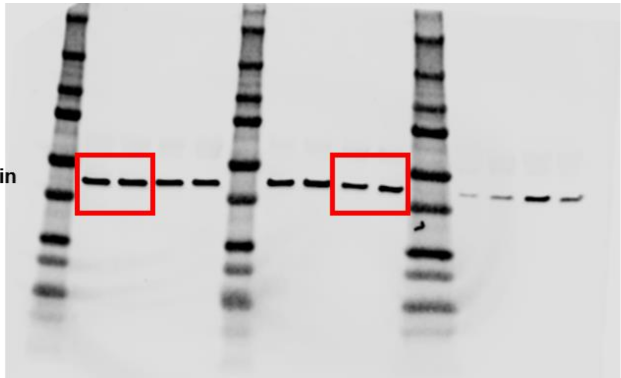

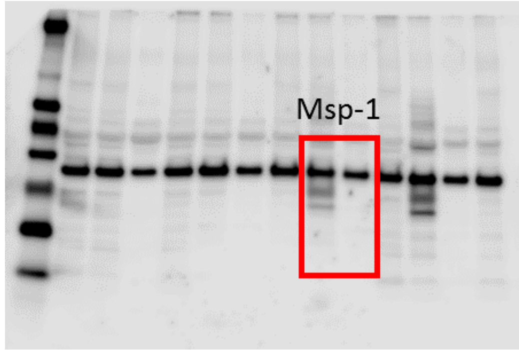


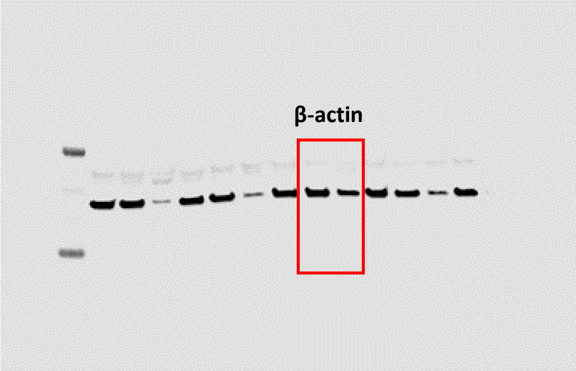


#### Supplementary Figure 1A

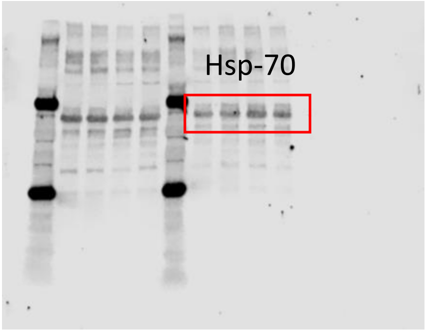


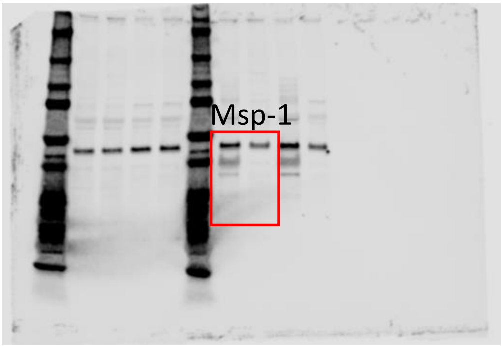

### Uncropped blots for Fig 1D

**β-actin**

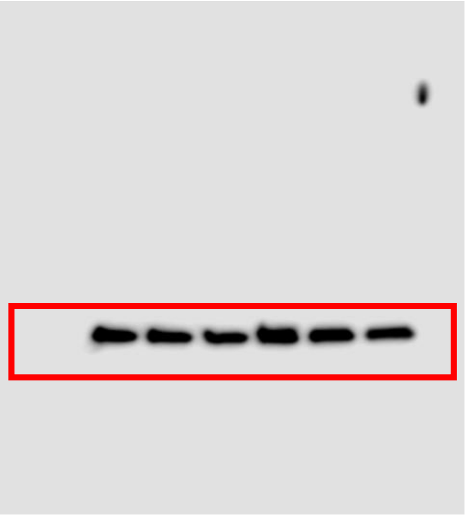

**p-p53$^{Ser15}$**

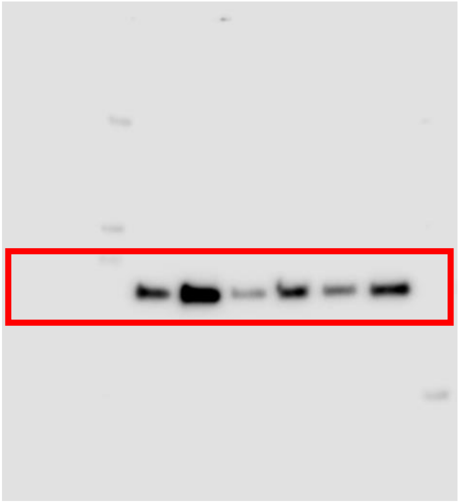

**p53**

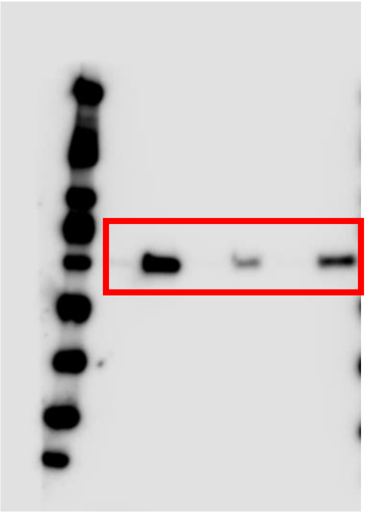

Fig2C

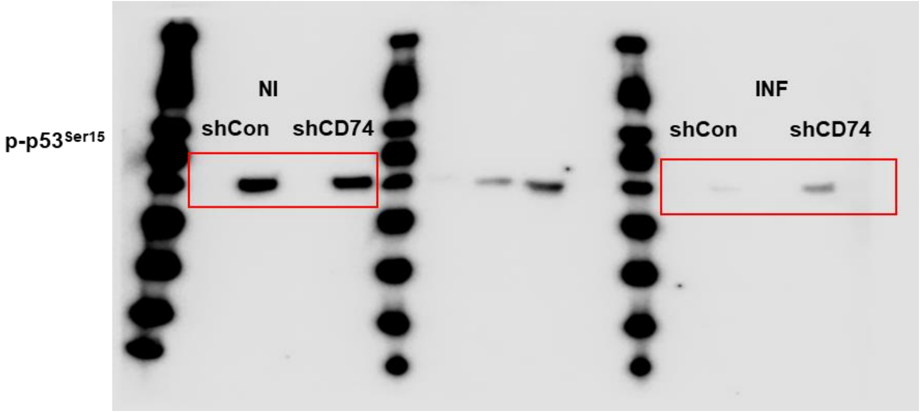
p-p53Ser15
NI
shCon
shCD74
INF
shCon
shCD74

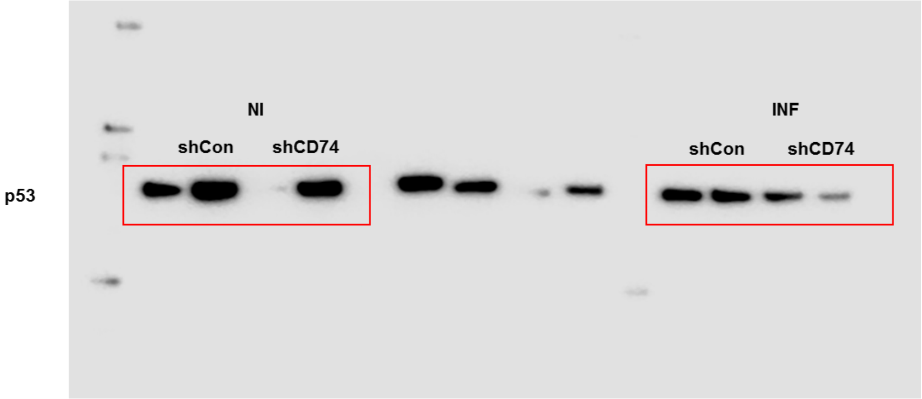
p53
NI
shCon
shCD74
INF
shCon
shCD74

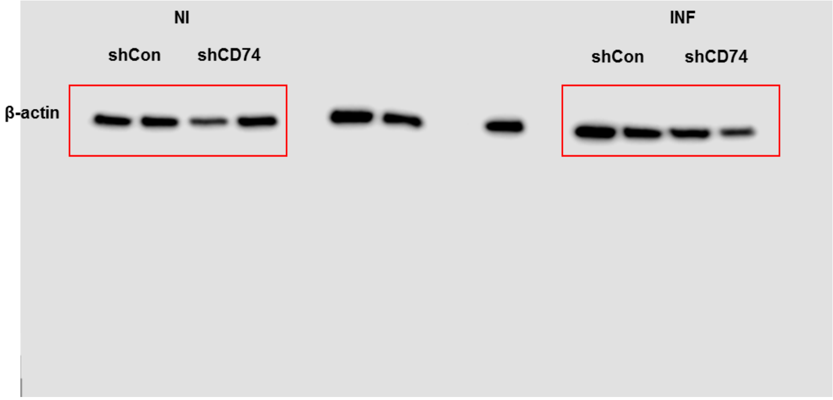
β-actin
NI
shCon
shCD74
INF
shCon
shCD74

Figure 3C

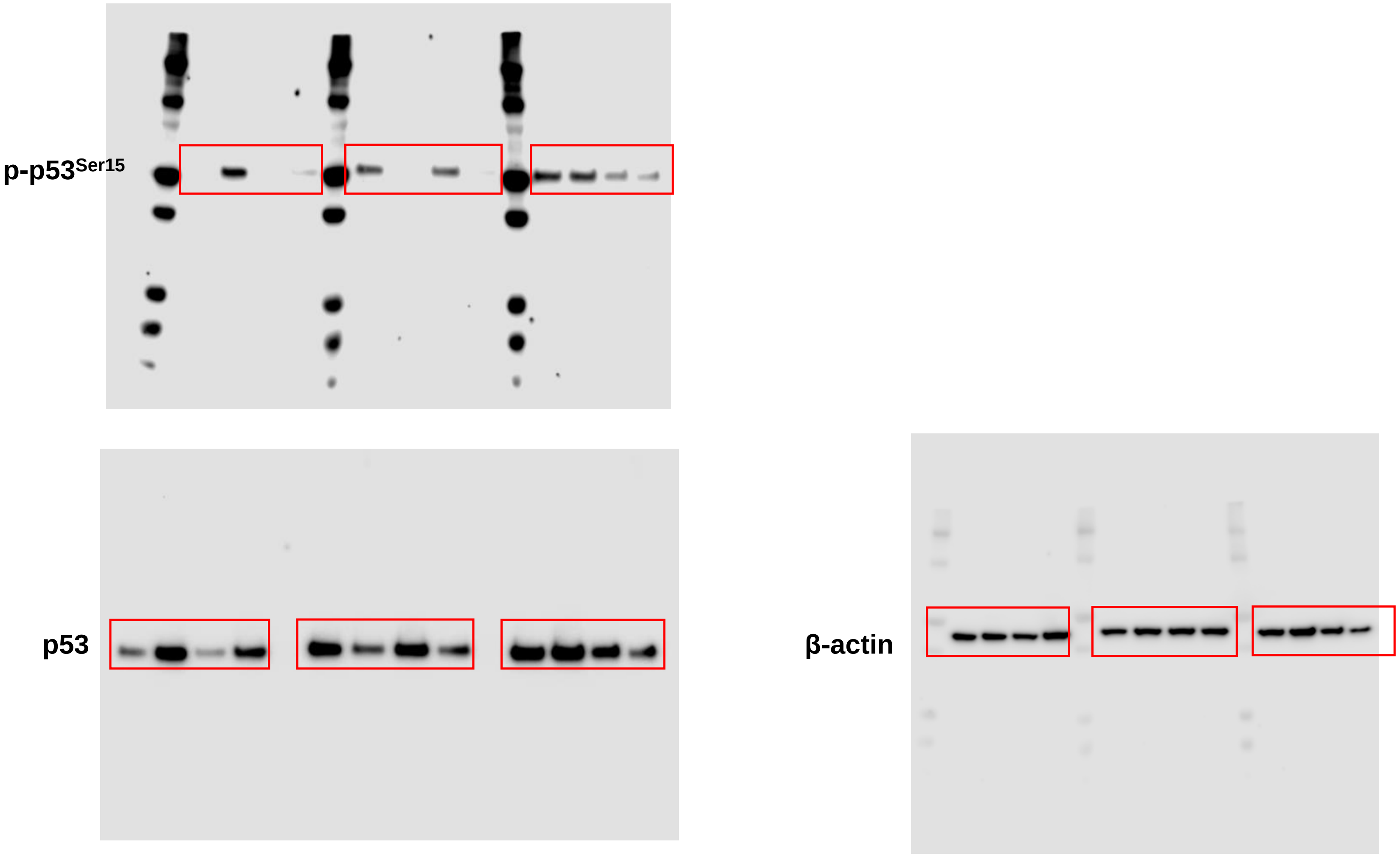

Supplementary Figure 1B

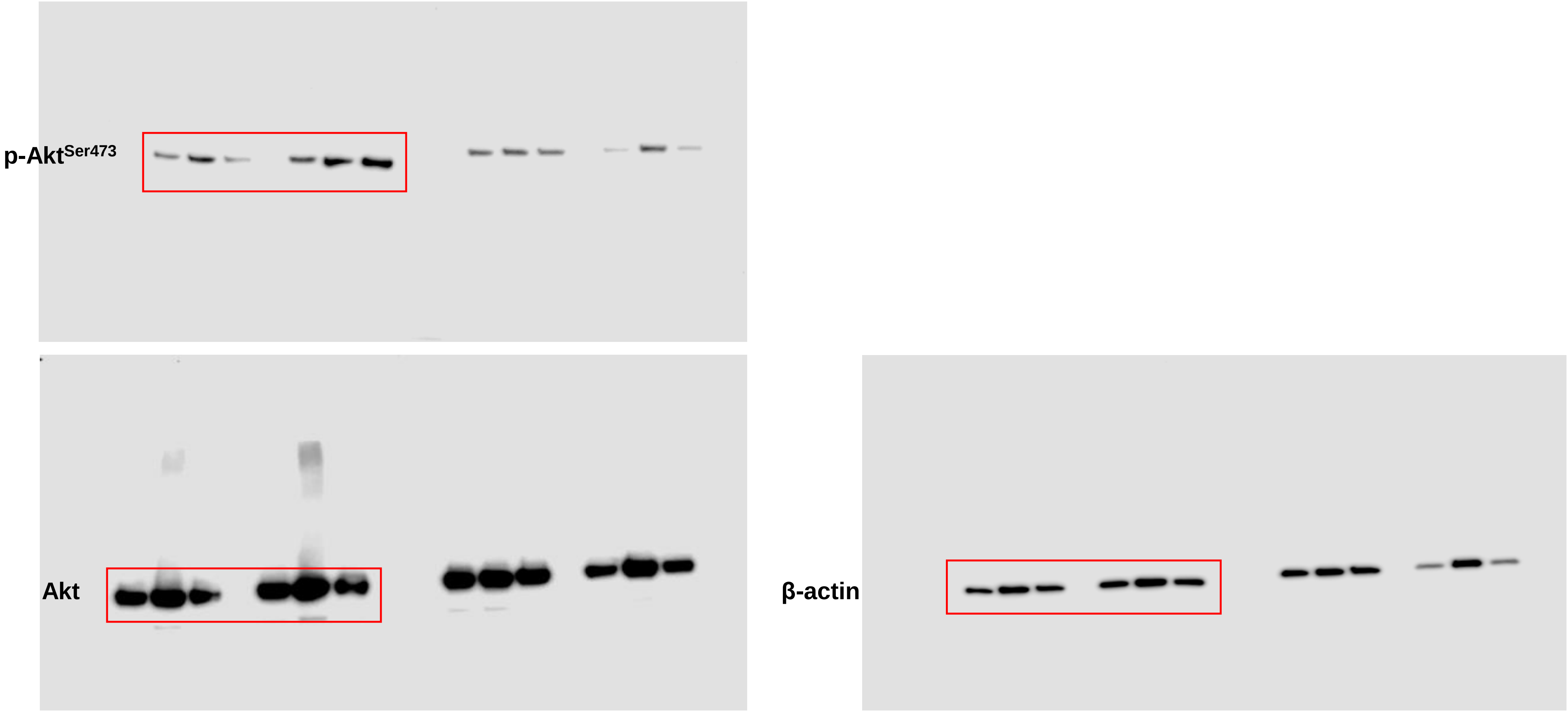
